# Multi-ethnic analysis shows genetic risk and environmental predictors interact to influence 25(OH)D concentration and optimal vitamin D intake

**DOI:** 10.1101/652941

**Authors:** Kathryn E. Hatchell, Qiongshi Lu, Julie A. Mares, Erin D. Michos, Alexis C. Wood, Corinne D. Engelman

## Abstract

**Background:** Vitamin D inadequacy affects almost 50% of adults in the United States and is associated with numerous adverse health effects. Vitamin D concentration [25(OH)D] is a complex trait with genetic and environmental predictors that work in tandem to influence 25(OH)D and may determine how much vitamin D intake is required to reach an optimal 25(OH)D concentration. To date, there has been little investigation into how genetics and environment interact to affect 25(OH)D.

**Objective:** Interactions between continuous measures of a polygenic score (PGS) and vitamin D intake (PGS*intake) or available ultra-violet (UV) radiation (PGS*UV) were evaluated separately in individuals of African or European ancestry.

**Methods:** Mega-analyses were performed using three independent cohorts (N=9,668; African ancestry n=1,099; European ancestry n=8,569). Interaction terms and joint effects (main and interaction terms) were tested using one-degree of freedom (DF) and 2-DF models, respectively. All models controlled for age, sex, body mass index (BMI), cohort, and dietary intake/available UV. Additionally, in participants achieving Institute of Medicine (IOM) vitamin D intake recommendations, 25(OH)D was evaluated by level of genetic risk of 25(OH)D deficiency.

**Results:** The 2-DF PGS*intake, 1-DF PGS*UV and 2-DF PGS*UV results were statistically significant in participants of European ancestry (p=3.3×10^−18^, 2.1×10^−2^, and 2.4×10^−19^, respectively), but not in those of African ancestry. In European-ancestry participants who reached IOM vitamin D intake guidelines, the percent of participants achieving adequate 25(OH)D (≥20ng/ml) increased as genetic risk decreased (72% vs 89% in the highest vs lowest risk categories; p=0.018).

**Conclusions:** Available UV radiation and vitamin D intake interact with genetics to influence 25(OH)D. Individuals with higher genetic risk of deficiency may require more vitamin D exposure to maintain optimal 25(OH)D concentrations. Overall, the results showcase the importance of incorporating both environmental and genetic factors into analyses, as well as the potential for gene-environment interactions to inform personalized dosing of vitamin D.

**Sources of Support:** *ARIC:* The Atherosclerosis Risk in Communities Study is carried out as a collaborative study supported by National Heart, Lung, and Blood Institute contracts (HHSN268201100005C, HHSN268201100006C, HHSN268201100007C, HHSN268201100008C, HHSN268201100009C, HHSN268201100010C, HHSN268201100011C, and HHSN268201100012C). The authors thank the staff and participants of the ARIC study for their important contributions. Funding for GENEVA was provided by National Human Genome Research Institute grant U01HG004402 (E. Boerwinkle).

*MESA:* MESA and the MESA SHARe project are conducted and supported by the National Heart, Lung, and Blood Institute (NHLBI) in collaboration with MESA investigators. Support for MESA is provided by contracts HHSN268201500003I, N01-HC-95159, N01-HC-95160, N01-HC-95161, N01-HC-95162, N01-HC-95163, N01-HC-95164, N01-HC-95165, N01-HC-95166, N01-HC-95167, N01-HC-95168, N01-HC-95169, UL1-TR-000040, UL1-TR-001079, UL1-TR-001420, UL1-TR-001881, and DK063491. The MESA CARe data used for the analyses described in this manuscript were obtained through Genetics (accession numbers). Funding for CARe genotyping was provided by NHLBI Contract N01-HC-65226. Funding support for the Vitamin D dataset was provided by grant HL096875

*WHI:* The WHI program is funded by the National Heart, Lung, and Blood Institute, National Institutes of Health, U.S. Department of Health and Human Services through contracts HHSN268201600018C, HHSN268201600001C, HHSN268201600002C, HHSN268201600003C, and HHSN268201600004C. This manuscript was not prepared in collaboration with investigators of the WHI, has not been reviewed and/or approved by the Women’s Health Initiative (WHI), and does not necessarily reflect the opinions of the WHI investigators or the NHLBI. WHI PAGE is funded through the NHGRI Population Architecture Using Genomics and Epidemiology (PAGE) network (Grant Number U01 HG004790). Assistance with phenotype harmonization, SNP selection, data cleaning, meta-analyses, data management and dissemination, and general study coordination, was provided by the PAGE Coordinating Center (U01HG004801-01). Funding support for WHI GARNET was provided through the NHGRI Genomics and Randomized Trials Network (GARNET) (Grant Number U01 HG005152). Assistance with phenotype harmonization and genotype cleaning, as well as with general study coordination, was provided by the GARNET Coordinating Center (U01 HG005157). Assistance with data cleaning was provided by the National Center for Biotechnology Information. Funding support for genotyping, which was performed at the Broad Institute of MIT and Harvard, was provided by the NIH Genes, Environment and Health Initiative [GEI] (U01 HG004424). The datasets used for the analyses described in this manuscript were obtained from dbGaP at http://www.ncbi.nlm.nih.gov/sites/entrez?db=gap through dbGa Paccession phs000200.v11.p3. Funding for WHI SHARe genotyping was provided by NHLBI Contract N02-HL-64278.

*Other:* KEH was supported by an NLM training grant to the Computation and Informatics in Biology and Medicine Training Program (NLM 5T15LM007359). Computational resources were supported by a core grant to the Center for Demography and Ecology at the University of Wisconsin-Madison (P2C HD047873). JM was supported by the Department of Ophthalmology and Visual Sciences, and by an unrestricted grant to the Department of Ophthalmology and Visual Sciences from the Research to Prevent Blindness, and by National Institutes of Health, National Eye Institute grants R01 EY016686, R01 EY025292.

## Introduction

Vitamin D inadequacy, as defined by a 25-hydroxyvitamin D [25(OH)D] concentration less than 20 ng/mL, affects almost 50% of adults in the United States (1-3). Low vitamin D concentrations have been associated with increased risk of autoimmune diseases, migraines, hypertension, dyslipidemia, cardiovascular events, and cardiovascular mortality (1, 3-9). Additionally, recent Mendelian randomization studies have suggested a causal relationship between low 25(OH)D concentrations and increased risk of obesity, ovarian cancer, hypertension, lower cognitive function in aging, multiple sclerosis, and all cause and cancer mortality (10-16). However, there have also been Mendelian randomization studies that found null relationships between vitamin D and coronary artery disease, depression and fatigue (17-19). Recent results from the Vitamin D and Omega-3 trial (VITAL) showed null associations between vitamin D supplementation and both cancer and cardiovascular disease. However, the inclusion of individuals with adequate 25(OH)D concentrations, lack of assessing individual 25(OH)D response to supplementation, and outside use of vitamin D before and during the trial limit the interpretability of these findings (20).

Serum 25(OH)D concentration is a complex phenotype with genetic and environmental predictors that may determine how much vitamin D intake is required to reach an optimal vitamin D blood concentration (21-24). Primary environmental predictors of 25(OH)D concentrations are vitamin D intake through diet and supplementation, and available ultraviolet (UV) radiation exposure. Therefore, knowledge of how genetic determinants of vitamin D concentrations interact with environmental predictors could be useful in the prevention of vitamin D associated morbidity and mortality. Understanding gene-by-environment interactions and how they affect 25(OH)D concentrations could inform personalized supplementation for maintaining adequate vitamin D concentrations through a precision public health approach.

While attention has been paid to genetic determinants of 25(OH)D concentration through genome-wide association studies (GWAS) and, separately, to the environmental determinants, much less research has focused on how environmental factors interact with genetic factors. Investigating the effects of genetic or environmental predictors in isolation may miss much of the variance in 25(OH)D. Through GWAS top findings, only 2.8% of the variance in 25(OH)D can be explained (25). Research has found that vitamin D intake through diet and supplement use accounts for 1-8% of the variation in vitamin D concentrations between individuals, and that sun exposure accounts for 1-15% of the variation (23, 26-28). One study in European-ancestry women reported an interaction between two SNPs in the GC gene (the GC protein product transports the vitamin D metabolites in the blood) and both vitamin D intake and sun exposure (23). The genetic effect was stronger, with more variance explained, in summer and in those with a higher intake of vitamin D. This same study reported preliminary evidence of differing genetic effect of a polygenic score (PGS) for deficiency, comprised of two SNPs, by level of vitamin D intake and season (23). Therefore, it is important to investigate gene-environment interactions as the risk inferred by genetic or environmental factors alone is not enough to predict risk of inadequate vitamin D concentrations.

To address these knowledge gaps, we examined the interactions between a new PGS and vitamin D intake or available UV radiation using separate linear models in individuals of African or European ancestry (29). Additionally, to replicate findings from a previous study (23), we also determined, using an independent cohort of participants achieving Institute of Medicine (IOM) vitamin D intake guidelines, the percent reaching adequate (>20 ng/ml) 25(OH)D concentrations, stratified by level of genetic risk. IOM vitamin D intake guidelines are 600 IU/day for those 70 years old or younger, and 800 IU for those over 70 years old. We hypothesized that gene*environment interactions would determine 25(OH)D concentrations. These results could help inform screening and treatment of vitamin D inadequacy based on genetic and environmental factors.

## Methods

### Participants

Analyses were performed in a sample of 8,569 participants of primarily European ancestry and 1,099 participants of primarily African ancestry who had data for the required variables: age, sex, body mass index (BMI), dietary intake of vitamin D, available UV radiation and genome-wide single nucleotide polymorphisms (SNPs). Participants were from Atherosclerosis Risk in Communities (ARIC), the Multi-Ethnic Study of Atherosclerosis (MESA) and the Women’s Health Initiative (WHI), and are independent of the GWAS meta-analysis, <u>TRANS</u>-ethni<u>C</u> <u>E</u>valuation of vitami<u>N</u> <u>D</u> (TRANSCEN-D), that provided the summary statistics used to calculate the PGS (30).

ARIC is a prospective study of men and women who were recruited from 4 U.S. locations [Forsyth County, NC; Jackson, MS; suburbs of Minneapolis, MN; and Washington County, MD] and were aged 46-70 years at visit 2 (1990-1992), which was the visit that serum 25(OH) was measured as part of an ancillary study. ARIC data were obtained through dbGaP Study Accession: phs000090.v4.p1. MESA is a prospective study of men and women aged 45-84 at baseline who were recruited from six United States sites: Columbia University, New York, NY; Johns Hopkins University, Baltimore, MD; Northwestern University, Chicago, IL; University of Minnesota, Minneapolis, MN; University of California at Los Angeles, Los Angeles, CA, and Wake Forest University, Winston-Salem, NC. Serum 25(OH)D was measured at MESA exam 1 (July 2000-August 2002). MESA data were obtained through dbGaP Study Accession: phs000209.v13.p3. Women participating in WHI were recruited from 40 clinical centers in the United States. Serum 25(OH)D was measured as part of the Calcium and Vitamin D (CaD) Trial (1993-1999) (31). WHI data were obtained through dbGaP Study Accession: phs000200.v11.p3. The data used in these analyses were collected under guidelines from the relevant institutional review boards and all participants provided informed consent, including consent for use of genetic data.

### Calculation of available UV radiation

Participants were assigned continuous available UV radiation values that were calculated based on month of blood draw and location using UV data from the National Weather Service Climate Prediction Center historical database. The UV radiation values ranged from 0.7 to 9.5 UV index units. The methods are described in more detail elsewhere (29).

#### Measurement of vitamin D intake

Dietary data were derived from study specific nutritional questionnaires; each study created a derived variable of typical dietary vitamin D intake. WHI also collected data on vitamin D supplement use at the same visit that 25(OH)D was assessed. The sum of vitamin D intake from food and supplements was derived and used for supplemental and sensitivity analyses, otherwise dietary intake alone was used. The methods are described in more detail elsewhere (29).

### Measurement of 25(OH)D

Serum 25(OH)D concentrations were measured by the studies using different assays, which are reported on elsewhere (29). To control for assay-level differences in vitamin D concentrations, vitamin D concentrations were converted to z-scores within studies for combined analyses.

### Data Quality Control

Quality control (QC) of phenotypic data is described in detail elsewhere (29). Generally, QC included winsorizing 25(OH)D to minimize the influence of outliers and using a log transformation to improve normality of 25(OH)D distribution in each cohort (32).

Where available for the respective visit, physical activity was measured in metabolic equivalent (MET) hours per week. Physical activity was capped at 16 MET hours/day or 112 MET hours/week. Additionally, physical activity data were normalized by cohort to account for the different surveys utilized to acquire the data.

### Genotyping and PGS Development

Genotyping methods are described in publications by ARIC, MESA, and WHI (33-37). Supplemental Table 1 gives information on the genotyping array used by the studies. QC was done in an ancestry-specific manner for those of European and African ancestry. QC methods are described in more detail elsewhere (29). **Supplemental Figures 1 and 2**, and **Supplemental Table 1** give specifics on quality control for each cohort. QC was performed using PLINK v1.9 and vcfTools (38, 39).

**Table 1:** Sample characteristics

| Cohort | Variable | European ancestry | African ancestry |
| --- | --- | --- | --- |
| ARIC | Sample size | 6,178 | 57 |
|  | Age (SE)<br>[years] | 57.1 (5.7) | 55.6 (6.2) |
|  | % Female | 54 | 49 |
|  | BMI (SE)<br>[kg/m<sup>2</sup>] | 27.3 (4.9) | 28.6 (5.7) |
|  | UV<sup>1</sup> (SE)<br>[units] | 5.1 (2.5) | 7.1 (2.4) |
|  | Intake<sup>2</sup> (SE)<br>[IU] | 223.3 (145.7) | 221.2 (137.3) |
|  | 25(OH)D (SE)<br>[ng/ml] | 26.0 (8.8) | 20.9 (7.8) |
| MESA | Sample size | 1,936 | 342 |
|  | Age (SE)<br>[years] | 62.7 (10.3) | 62.3 (10.4) |
|  | % Female | 53 | 51 |
|  | BMI (SE)<br>[kg/m<sup>2</sup>] | 27.8 (5.0) | 30.0 (5.9) |
|  | UV (SE)<br>[units] | 4.5 (2.3) | 5.1 (2.2) |
|  | Intake (SE)<br>[IU] | 188.9 (157.2) | 161.8 (144.1) |
|  | 25(OH)D (SE)<br>[ng/ml] | 30.1 (10.8) | 19.5 (8.9) |
| WHI | Sample size | 455 | 700 |
|  | Age (SE)<br>[years] | 66.6 (6.8) | 61.8 (7.4) |
|  | % Female | 100 | 100 |
|  | BMI (SE)<br>[kg/m<sup>2</sup>] | 29.9 (6.3) | 31.1 (6.4) |
|  | UV (SE)<br>[units] | 5.2 (2.5) | 5.5 (2.6) |
|  | Intake (SE)<br>[IU] | 192.3 (143.2) | 146.4 (130.5) |
|  | 25(OH)D (SE)<br>[ng/ml] | 18.9 (10.7) | 19.0 (15.4) |

<sup>1</sup>available UV radiation
<sup>2</sup>vitamin D intake from diet
<sup>3</sup>MANOVA global test (performed in SAS (version 9.4) revealed differences in one or more variables by cohort, therefore cohort was adjusted for in all models that included multiple cohorts

**Figure 1:**
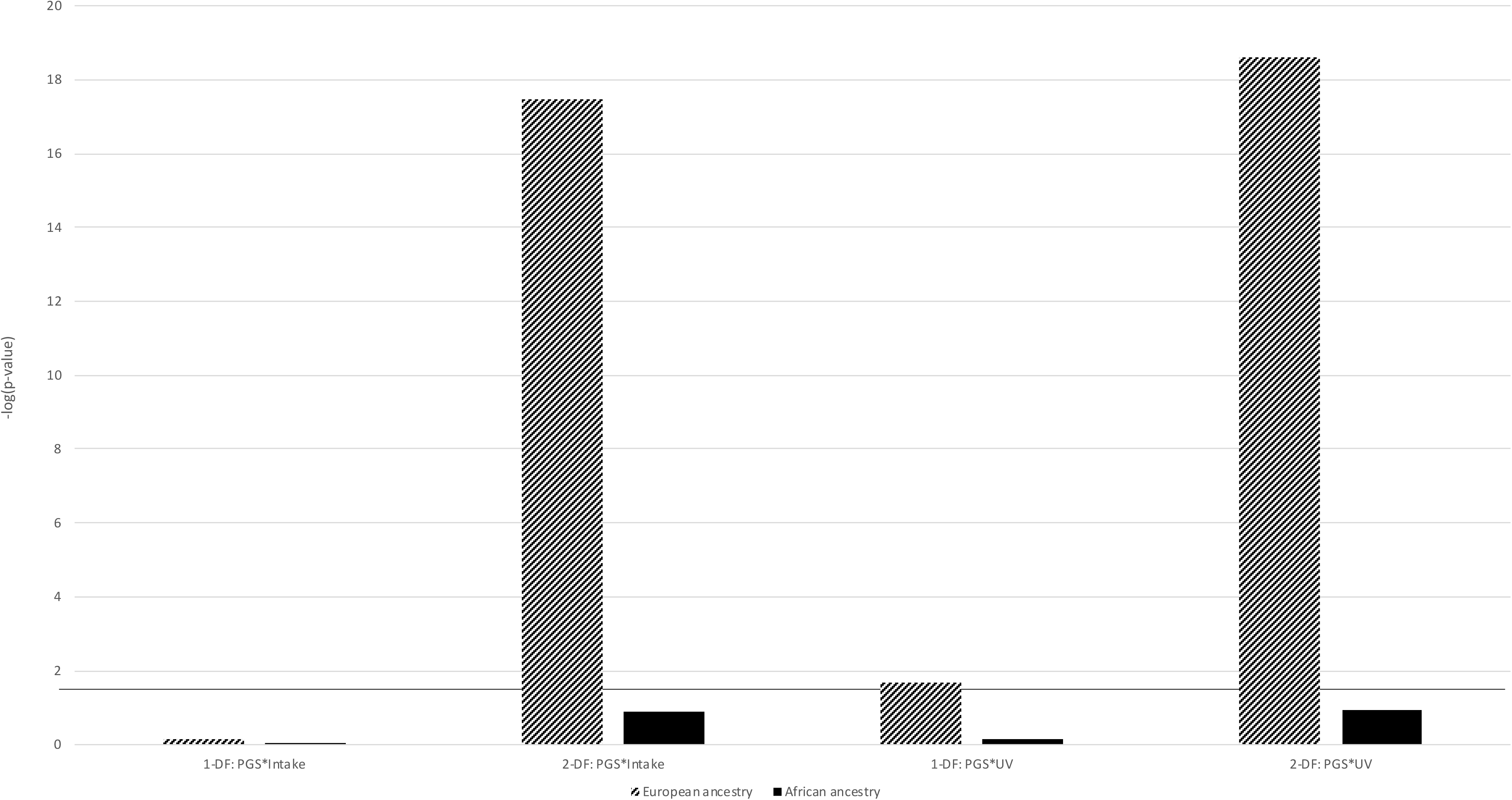
p-values for G*E interaction terms from 1-DF and 2-DF models. Figure 1 shows –log(p-values) for the 1-DF and 2-DF models of the PGS interaction term or joint effect, respectively; all models controlled for age, sex, BMI, cohort, vitamin D intake and available UV radiation. The black horizontal line denotes the p=0.05 significance cutoff. The 2-DF PGS*intake, 1-DF PGS*UV and 2-DF PGS*UV results were statistically significant in participants of European ancestry (p=3.3×10^−18^, 2.1×10-2 and 2.4×10^−19^, respectively).

**Figure 2:**
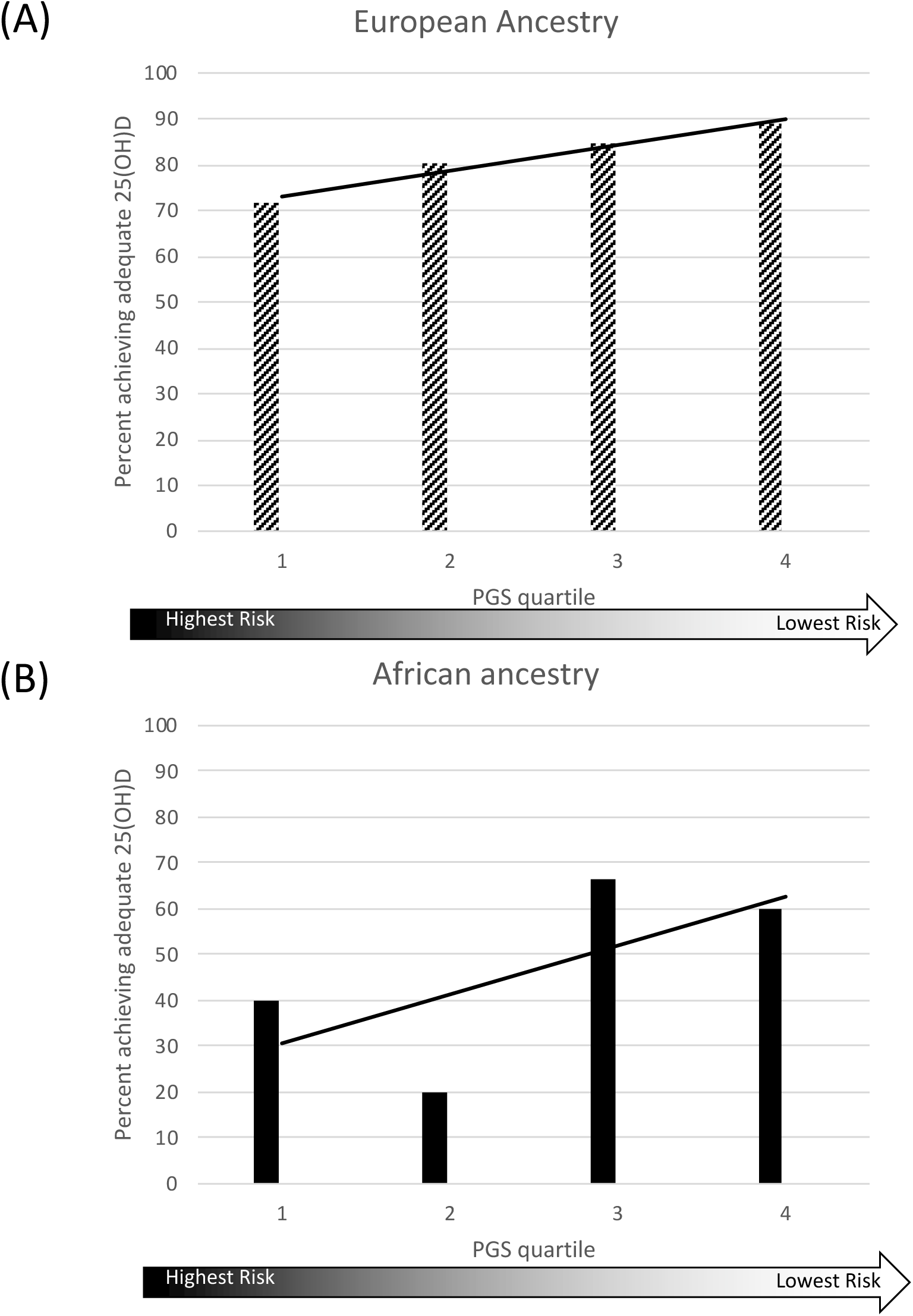
Percent achieving adequate 25(OH)D in those reaching IOM vitamin D intake guidelines by genetic risk. Figure 2 shows the percent of European- or African-ancestry participants who reached IOM vitamin D intake guidelines and achieved adequate 25(OH)D (20 ng/ml) by quartile of genetic risk. In those of European ancestry (panel (A), n=184), as genetic risk decreased (higher PGS), those reaching optimal vitamin D concentrations increased. The difference in percent reaching adequate 25(OH)D between the two extreme quartiles was 17.3%; 71.7% of participants with the highest genetic risk and 89.0% of participants with the lowest risk reached adequate 25(OH)D. This is a statistically significant (p=0.018) and clinically meaningful difference. The trend was not significant in those of African ancestry (panel (B), n=17).

Previously, an optimal PGS was determined in an ancestry-specific manner for those of European or African ancestries (29). PGSs were weighted using effect sizes from an independent multi-ethnic GWAS, TRANSCEN-D, the largest multi-ethnic vitamin D GWAS meta-analysis to date (40). European summary statistics came from the SUNLIGHT discovery cohort that was included in TRANSCEN-D. After QC, 8,569 European samples and 1,099 African-ancestry samples remained for the analysis.

### Statistical Analysis

Two separate sets of models were investigated: 1) one-degree of freedom (DF) models which tested only the relevant interaction term and 2) 2-DF models which jointly tested both the relevant interaction term and the PGS main effect term (41). Relevant interaction terms were the PGS interacting with either vitamin D intake (PGS*intake) or available UV radiation (PGS*UV). All 1-DF and 2-DF models controlled for age, sex, BMI, cohort, vitamin D intake, and available UV radiation. All statistical analyses were performed using SAS (version 9.4). Further analyses were performed in those who achieved IOM vitamin D intake guidelines (600 IU/day for those 1-70 years old and 800 IU/day for those over 70) to explore differences in the percent of those achieving adequate 25(OH)D concentrations (≥20 ng/ml) by decile of genetic risk. Statistical significance was determined by testing difference between two proportions.

Sensitivity analyses were performed in a subset of participants with physical activity data or vitamin D supplement use data, which permitted adjusting for these variables. Additionally, sensitivity analyses controlling for principal components (PCs) of ancestry were performed. All sensitivity analyses were performed in an ancestry-specific manner for European and African cohorts. Additional sensitivity analyses were performed to ensure that the randomized controlled trial (RCT) study design of the WHI CaD trial was not biasing the results.

## Results

Participant characteristics of this sample can be found in **Table 1.** Gene-environment interactions for PGS*intake and PGS*UV were tested for with a 1-DF and 2-DF approach (**Table 2 and Figure 1**). The interaction term in the 1-DF PGS*UV model and the joint effect in the 2-DF PGS*intake and 2-DF PGS*UV models were statistically significant in participants of European ancestry (p=2.1×10^−2^, 3.3×10^−18^ and 2.4×10^−19^, respectively). In African-ancestry analyses, power was limited due to the smaller sample size, and no statistically significant interactions or joint effects were discerned.

**Table 2:** Betas, standard errors and p-values for G*E interaction terms

| Model | European Ancestry |  |  |  | African Ancestry |  |  |  |
| --- | --- | --- | --- | --- | --- | --- | --- | --- |
|  | PGS*UV |  | PGS*Intake |  | PGS*UV |  | PGS*Intake |  |
|  | Beta (SE) | p-value | Beta (SE) | p-value | Beta (SE) | p-value | Beta (SE) | p-value |
| Environmental main effect<sup>1</sup> | 0.096<br>(0.005) | <0.0001 | 0.11<br>(0.011) | <0.0001 | 0.07<br>(0.012) | <0.0001 | 0.13<br>(0.033) | 0.0002 |
| Genetic main effect<sup>2</sup> | 0.087<br>(0.038) | 0.022 | 0.16<br>(0.018) | <0.0001 | 0.086<br>(0.071) | 0.23 | 0.062<br>(0.031) | 0.04 |
| Interaction term | 0.017<br>(0.0073) | 0.021 | 0.0006<br>(0.018) | 0.74 | -0.0044<br>(0.012) | 0.71 | 0.00042<br>(0.03) | .99 |

<sup>1</sup>Environmental main effect is either available UV radiation (UV) measured for month prior to blood draw (values ranged from 0.7 to 9.5 units), or vitamin D intake through diet (Intake) assessed at blood draw visit and measured in IU
<sup>2</sup>Genetic main effect term is the PGS
<sup>3</sup>All models controlled for age, sex, BMI, cohort and vitamin D intake or available UV radiation

Sensitivity analyses were performed. Characteristics for participants used in sensitivity analyses can be found in **Supplemental Tables 2 and 3**. Sensitivity analyses controlling for physical activity showed the same pattern of significance for interaction terms, however, p-values were slightly attenuated due to smaller sample size (**Supplemental Figure 3**). Interaction terms were no longer significant in the sensitivity analyses that used the subsample with vitamin D supplement use, due to loss of power and small sample size (European ancestry n=455; African ancestry n=700). Adding PCs to analyses had only a minor impact on effect size and did not change significance or interpretation. To ensure the RCT study design of WHI did not influence the results, additional sensitivity analyses were performed. There was no significant difference in 25(OH)D concentration between participants on the treatment arm compared to the placebo arm. Additionally, there was no significant difference in the association between the PGS and 25(OH)D in WHI compared to the other cohorts.

Next, in participants who reached IOM vitamin D dietary intake guidelines, the percent of participants achieving adequate 25(OH)D concentration by PGS quartile was calculated (**Figure 2**). In those of European ancestry, as genetic risk decreased, those reaching optimal vitamin D concentrations increased (71.7% vs 89.0% in the highest and lowest risk categories, respectively). This is a statistically significant (p=0.018) and clinically meaningful difference. The directionality of the trend persisted in those of African ancestry, however, the difference was not significant (p=0.28). This confirmed results previously reported on in an independent sample (23).

## Discussion

Findings presented here build upon existing literature reporting that UV radiation and vitamin D intake modify the effect that genetics have on 25(OH)D concentrations (23, 42). Previously, in women of European ancestry, the genetic effects were reported to be stronger in summer and in women with high vitamin D intake (>400 IU/day) (23). Here, these results are replicated in both women and men of European ancestry. As available UV radiation or dietary vitamin D intake increased, the genetic risk score had a larger effect on 25(OH)D (**Table 2**; interaction term not significant for dietary vitamin D intake). The current study also includes individuals of African ancestry, but results were not significant, likely a reflection of the smaller sample size and, subsequently, reduced power.

Gene-environment interactions can have public health implications, especially in the context of 25(OH)D. Extending previous findings in women of European ancestry (20), the results presented here in men and women of African or European ancestry indicate that, for those of high genetic risk, IOM recommendations for vitamin D intake may not be sufficient. As shown by about a third of those with highest genetic risk not reaching adequate 25(OH)D albeit achieving IOM recommendations for vitamin D intake. In European-ancestry participants who reached IOM dietary guidelines for vitamin D intake, significantly fewer participants with high genetic risk reached optimal 25(OH)D concentrations (≥20 ng/ml). This trend was also seen in African-ancestry participants, although it was not statistically significant, possibly due to the relatively small sample size. These results suggest that a precision public health approach to achieve adequate blood levels of vitamin D may be more effective, tailoring intake recommendations to genetic risk.

While this study builds upon the novel interactions previously reported (20) by including both men and women of European and African ancestry, it is not without limitations. First while exploring gene-by-environment interactions that influence 25(OH)D concentrations in a multi-ethnic sample is novel, the relatively small size of the African-ancestry sample limited the power. To maintain independence from TRANSCEN-D, which provided ancestry-specific weights for the PGSs, the sample size used in this analysis was relatively small, especially for the African-ancestry cohort (n=1,099), as nearly all the publicly available African-ancestry samples with relevant data had been exhausted. This emphasizes that we, as a research community, need to include more individuals of African ancestry in our studies to better understand vitamin D requirements and other health outcomes and make ancestrally informed recommendations that combat instead of accentuate health disparities (i.e. in initiatives like All of Us) (43, 44). Additionally, while the use of available UV radiation is a substantial improvement from using season as a measure of UV exposure, it does not include behaviors which can alter an individual’s exposure in months when UV exposure is high enough to make substantial vitamin D in the skin, such as the time spent outside, the amount of skin usually exposed, and sunblock use. The lack of information about these behaviors could introduce measurement error to the estimate of an individual’s UV radiation and, consequently, limit power. Finally, vitamin D supplementation is a stronger predictor of 25(OH)D concentrations than vitamin D intake from food, which is generally in much lower amounts than those found in supplements. However, only the WHI study measured vitamin D supplement intake for the relevant visit. Therefore, only interactions involving dietary intake had adequate sample size to be explored in this study, which could have led to the lack of a significant interaction being detected between the PGS and vitamin D intake in the 1-DF models. Nonetheless, findings here guide future research in the quest for precision public health management of 25(OH)D inadequacy.

## Conclusion

This research adds to the ongoing narrative deciphering the predictors of 25(OH)D concentrations, by extending evidence suggesting that levels of environmental sources of vitamin D (intake and UV radiation) affect 25(OH)D concentrations differently in those with low versus high genetic risk, reiterating the importance of well-measured environmental factors in genetic analyses. This research also adds to evidence indicating the importance of considering genetic risk when making recommendations on vitamin D intake and personalizing the dose of vitamin D to best achieve optimal 25(OH)D concentrations.

## Statement of authors’ contributions to manuscript

K.H. and C.E. designed research; K.H. conducted research; K.H. analyzed data; Q.L. provided statistical guidance; A.W. provided MESA vitamin D intake data; J.M. provided nutritional insight; and K.H. wrote the paper. K.H. and C.E. had primary responsibility for final content. E.D.M contributed intellectual content to paper. All authors read and approved the final manuscript.

