## Supplemental Material for "Multi-ethnic analysis shows genetic risk and environmental predictors interact to influence 25(OH)D concentration and optimal vitamin D intake"

**Supplementary Data**

###### Supplemental Table 1: Genotyping information and quality control by cohort

| **Cohort** | **Genotyping Array** | **n** | **SNP quality control** | | | | **Sample quality control** | |
| --- | --- | --- | --- | --- | --- | --- | --- | --- |
|  |  |  | **Call rate** | **Minor Allele Frequency** | **Hardy-Weinberg Equilibrium (post-imputation)** | **# SNPs Passing QC** | **Call rate** | **Exclusions** |
| ARIC | AffymetrixGenome-WideHuman SNPArray 6.0 | African ancestry: 1,908  European ancestry: 7,178 | 95% | 0.002 | African Ancestry: 6x10^-8^ (5x10^-9^)    European Ancestry: 6x10^-8^ (6x10^-9^) | African ancestry: 9,335,785  European ancestry: 8,315,761 | 95% | sex mismatch, relatedness, chromosomal abnormalities |
| MESA | Affymetrix 50K gene-focused molecular imprinted polymer array (CVDSNP55v1_A) | African ancestry: 1,176  European ancestry: 1,936 | 95% | 0.002 | African Ancestry: 1x10^-6^ (2x10^-7^)  European Ancestry: 1x10^-6^ (1x10^-7^) | African ancestry: 309,712  European ancestry: 455,155 | 95% | sex mismatch, relatedness |
| WHI (Child6 NHLBI cohort) | AffymetrixGenome-WideHuman SNPArray 6.0 | African ancestry  Consent group 1: 65  Consent group 2: 572 | 95% | 0.002 | Consent group 1: 6x10^-8^ (5x10^-9^)  Consent group 2: 6x10^-8^ (5x10^-9^) | Consent group 1: 9,551,098  Consent group 2: 9,997,380 | 95% | sex mismatch, relatedness, race mismatch |
| WHI (Child7 GARNET cohort) | Illumina HumanOmni1-Quad v1-0 B | European ancestry:  Consent group 1: 86  Consent group 2: 443 | 95% | 0.002 | Consent group 1: 5x10^-8^ (5x10^-9^)  Consent group 2: 5x10^-8^ (4x10^-9^) | Consent group 1: 9,2037,621  Consent group 2: 9,722,526 | 95% | sex mismatch, relatedness, chromosome anomalies, race mismatch |
| WHI (Child 9 PAGE cohort) | Illumina MEGA Consortium 15063755 B2 array | African ancestry: 63 | 95% | 0.002 | 3x10^-8^ (5x10^-9^) | African ancestry: 9,641,566 | 95% | sex mismatch, relatedness, chromosome anomalies, race mismatch |

^a^All African-ancestry samples were imputed to CAAPA using Michigan Imputation Server

^b^All European-ancestry samples were imputed to HRC r1.1 2016 using Michigan Imputation Server

^c^Sample sizes reported here reflect entire eligible cohort, actual numbers used differ to maintain independence where necessary

###### Supplemental Figure 1**:** Pre-imputation quality-control process

- ARIC:
  - European American: 1,914 participants remain
  - African American: 7,197 participants remain
- MESA:
  - European American: 1,176 participants remain
  - African American: 1,936 participants remain
- WHI
  - NHLBI consent group 1 (African American): 65 participants remain
  - GARNET consent group 1 (European American): 87 participants remain
  - PAGE consent group 1 (African American): 118 participants remain
  - NHLBI consent group 2 (African American): 572 participants remain
  - GARNET consent group 2 (European American): 443 participants remain
- ARIC:
  - European American: 1,915 participants remain
  - African American: 7,197 participants remain
- MESA:
  - European American: 1,176 participants remain
  - African American: 1,940 participants remain
- WHI
  - NHLBI consent group 1 (African American): 66 participants remain
  - GARNET consent group 1 (European American): 87 participants remain
  - PAGE consent group 1 (African American): 118 participants remain
  - NHLBI consent group 2 (African American): 572 participants remain
  - GARNET consent group 2 (European American): 443 participants remain
- ARIC:
  - European American: 798,869 SNPs remain
  - African American: 720,277 SNPs remain
- MESA:
  - European American: 43,755 SNPs remain
  - African American: 36,900 SNPs remain
- WHI
  - NHLBI consent group 1 (African American): 853,490 SNPs remain
  - GARNET consent group 1 (European American): 804,152 SNPs remain
  - PAGE consent group 1 (African American): 1,125,184 SNPs remain
  - NHLBI consent group 2 (African American): 861,780 SNPs remain
  - GARNET consent group 2 (European American): 812,797 SNPs remain
- ARIC:
  - European American: 802,392 SNPs remain
  - African American: 799,260 SNPs remain
- MESA:
  - European American: 46,303 SNPs remain
  - African American: 46,319 SNPs remain
- WHI
  - NHLBI consent group 1 (African American): 863,945 SNPs remain
  - GARNET consent group 1 (European American): 954,133 SNPs remain
  - PAGE consent group 1 (African American): 1,479,901 SNPs remain
  - NHLBI consent group 2 (African American): 865,600 SNPs remain
  - GARNET consent group 2 (European American): 954,463 SNPs remain
- ARIC: 805,099 SNPs remain
- MESA: 46,425 SNPs remain
- WHI
  - NHLBI consent group 1 (African American): 863,948 SNPs remain
  - GARNET consent group 1 (European American): 945,139 SNPs remain
  - PAGE consent group 1 (African American): 1,481,282 SNPs remain
  - NHLBI consent group 2 (African American): 866,729 SNPs remain
  - GARNET consent group 2 (European American): 954,544 SNPs remain
- ARIC: 9,112 participants remain
- MESA: 3,116 participants remain
- WHI
  - NHLBI consent group 1 (African American): 66 participants remain
  - GARNET consent group 1 (European American): 87 participants remain
  - PAGE consent group 1 (African American): 118 participants remain
  - NHLBI consent group 2 (African American): 577 participants remain
  - GARNET consent group 2 (European American): 449 participants remain

Remove samples with call rate <95%

Remove SNPs with call rate <95%

Remove SNPs not in HWE

Remove SNPs with MAF <0.002

Remove one participant from pairs with IBD >0.38

Remove racial/ethnic mismatches

Supplemental Figure 1 shows pre-imputation quality control steps and cutoffs used as well as corresponding sample sizes at each step.

- ARIC:
  - African American: 1,914 participants remain
  - European American: 7,197 participants remain
- MESA:
  - African American: 1,179 participants remain
  - European American: 1,936 participants remain
- WHI
  - NHLBI consent group 1 (African American): 65 participants remain
  - GARNET consent group 1 (European American): 87 participants remain
  - PAGE consent group 1 (African American): 118 participants remain
  - NHLBI consent group 2 (African American): 572 participants remain
  - GARNET consent group 2 (European American): 443 participants remain

###### Supplemental Figure 2: Quality-Control process starting at imputation

- ARIC:
  - European American: 71,425 SNPs removed
  - African American: 100,921 SNPs removed
- MESA:
  - European American: 3,208 SNPs removed
  - African American: 5,189 SNPs removed
- WHI
  - NHLBI consent group 1 (African American): 107,984 SNPs removed
  - GARNET consent group 1 (European American): 44,890 SNPs removed
  - PAGE consent group 1 (African American): 124,466 SNPs removed
  - NHLBI consent group 2 (African American): 109,381 SNPs removed
  - GARNET consent group 2 (European American): 46,271 SNPs removed
- ARIC:
  - European American: 9,515,145 SNPs remain
  - African American: 8,931,534 SNPs remain
- MESA:
  - European American: 313,999 SNPs remain
  - African American: 478,047 SNPs remain
- WHI
  - NHLBI consent group 1 (African American): 9,540,068 SNPs remain
  - GARNET consent group 1 (European American): 9,134,809 SNPs remain
  - PAGE consent group 1 (African American): 9,588,806 SNPs remain
  - NHLBI consent group 2 (African American): 10,158,175 SNPs remain
  - GARNET consent group 2 (European American): 11,399,322 SNPs remain
- ARIC:
  - European American: 9,524,430 SNPs remain
  - African American: 8,933,380 SNPs remain
- MESA:
  - European American: 314,506 SNPs remain
  - African American: 477,815 (removed multi-allelic SNPs) SNPs remain
- WHI
  - NHLBI consent group 1 (African American): 9,551,099 SNPs remain
  - GARNET consent group 1 (European American): 9,126,383 remain
  - PAGE consent group 1 (African American): 9,657,908 SNPs remain
  - NHLBI consent group 2 (African American): 10,168,545 SNPs remain
  - GARNET consent group 2 (European American): 11,399,322 SNPs remain
- ARIC:
  - European American: 1,908 participants and 9,335,786 SNPs remain
  - African American: 7,178 participants and 8,315,761 SNPs remain
- MESA:
  - European American: 1,179 participants and 309,712 SNPs remain
  - African American: 1,936 participants and 455,155 SNPs remain
- WHI
  - NHLBI consent group 1 (African American): 65 participants and 9,551,098 SNPs remain
  - GARNET consent group 1 (European American): 86 participants and 9,087,621 remain
  - PAGE consent group 1 (African American): 63 participants and 9,641,566 SNPs remain (45 participants were duplicates to NHLBI consent group 2 and were removed)
  - NHLBI consent group 2 (African American): 572 participants and 9,997,380 SNPs remain
  - GARNET consent group 2 (European American): 443 participants and 9,722,526 SNPs remain
- ARIC:
  - European American: 7,178 participants and 39,172,678 SNPs remain [22 participants removed – fell below call rate >95% with removals of SNPs on wrong strand)
  - African American: 1,908 participants and 29,839,436 SNPs remain (6 participants removed – fell below call rate >95% with removals of SNPs on wrong strand)
- MESA:
  - European American: 1,936 participants and 39,072,978 SNPs remain
  - African American: 1,176 participants and 29,805,153 participants remain
- WHI
  - NHLBI consent group 1 (African American): 65 participants and 229,840,263 SNPs remain
  - GARNET consent group 1 (European American): 86 participants and 39,127,678 SNPs remain
  - PAGE consent group 1 (African American): 108 participants and 28,983,707 SNPs remain (1 subject removed – fell below call rate <95% with removals of SNPs on wrong strand)
  - NHLBI consent group 2 (African American): 572 participants and 29,840,474 SNPs remain
  - GARNET consent group 2 (European American): 443 participants and 39,127,678 SNPS remain

Remove SNPs on the wrong strand

Impute samples with Michigan Imputation Server

Remove SNPs with quality score <0.08 or MAF <0.001

Merge original genotypes back in

Final QC (call rate, HWE and MAF)

Supplemental Figure 2 shows quality control steps and cutoffs used as well as corresponding sample sizes at each step starting at the imputation phase. Sample sizes reported here reflect entire eligible cohort, actual numbers used differ to maintain independence where necessary.

###### Supplemental Table 2: Characteristics of sub-sample with supplement use data

| **Cohort** | **Variable** | **European-ancestry** | **African-ancestry** |
| --- | --- | --- | --- |
| **WHI** | **Sample size** | 455 | 700 |
|  | **Age (SE) [years]** | 66.6 (6.8) | 61.8 (7.4) |
|  | **% Female** | 100 | 100 |
|  | **BMI (SE) [kg/m^2^]** | 29.9 (6.3) | 31.2 (6.4) |
|  | **UV (SE)^1^ [units]** | 5.2 (2.5) | 5.5 (2.6) |
|  | **Intake (SE)^2^ [IU]** | 420.9 (299.4) | 308.8 (257.4) |
|  | **25(OH)D (SE) [ng/ml]** | 18.9 (10.7) | 19.0 (15.4) |

^1^available UV radiation
^2^vitamin D intake from diet and supplements

###### Supplemental Table 3: Characteristics of sub-sample with physical activity data

| **Cohort** | **Variable** | **European-ancestry** | **African-ancestry** |
| --- | --- | --- | --- |
| **MESA** | **Sample size** | 1,935 | 341 |
|  | **Age (SE) [years]** | 62.7 (10.3) | 62.3 (10.4) |
|  | **% Female** | 53% | 52% |
|  | **BMI (SE) [kg/m^2^]** | 27.8 (5.0) | 30.1 (5.9) |
|  | **UV (SE)^1^ [units]** | 4.5 (2.3) | 161.4 (144.1) |
|  | **Intake (SE)^2^ [IU]** | 188.9 (157.2) | 25.7 (32.0) |
|  | **Met-hours/week (SE)** | 26.0 (28.2) | 24.8 (28.8) |
|  | **25(OH)D (SE) [ng/ml]** | 30.1 (10.9) | 19.5 (8.9) |
| **WHI** | **Sample size** | 436 | 363 |
|  | **Age (SE) [years]** | 66.6 (6.8) | 61.9 (7.6) |
|  | **% Female** | 100 | 100 |
|  | **BMI (SE) [kg/m^2^]** | 29.9 (6.3) | 31.9 (6.4) |
|  | **UV (SE) [units]** | 5.2 (2.5) | 5.5 (2.6) |
|  | **Intake (SE) [IU]** | 193.6 (144.7) | 145.0 (119.4) |
|  | **Met-hours/week (SE)** | 6.3 (9.5) | 6.2 (11.0) |
|  | **25(OH)D (SE) [ng/ml]** | 18.9 (10.8) | 16.8 (12.9) |

^1^available UV radiation

^2^vitamin D intake from diet

###### Supplemental Figure 3: Sensitivity analyses interaction test results from 1-DF and 2-DF models controlling for physical activity

-log(p-value)

|  | European ancestry | African ancestry |
| --- | --- | --- |
| 1-DF: PGS*Intake | 0.05061 | 1.06248 |
| 2-DF: PGS*Intake | 15.4949 | 0.44406 |
| 1-DF: PGS*UV | 0.552842 | 0.0015227 |
| 2-DF: PGS*UV | 15.7447 | 0.000608438 |

Supplemental Figure 3 shows –log(p-values) for the 1-DF and 2-DF models of the PGS interaction term or joint effect; all models controlled for age, sex, BMI, cohort, physical activity, vitamin D intake and available UV radiation. The black horizontal line denotes the p=0.05 significance cutoff. The 2-DF PGS*intake and 2-DF PGS*UV results were statistically significant in participants of European ancestry (p=3.2x10^-16^and 1.8x10^-16^, respectively). Original to this manuscript.
